# Deep Learning Improves Parameter Estimation in Reinforcement Learning Models

**DOI:** 10.1101/2025.03.21.644663

**Authors:** Hua-Dong Xiong, Li Ji-An, Marcelo G. Mattar, Robert C. Wilson

**Affiliations:** School of Psychology, Georgia Institute of Technology; Neurosciences Graduate Program, University of California San Diego; Department of Psychology, New York University

**Keywords:** Cognitive modeling, Parameter estimation, Reinforcement learning, Decision making

## Abstract

Cognitive models are widely used in psychology and neuroscience to formulate and test hypotheses about cognitive processes. These processes are characterized by model parameters, which are then used for scientific inference. The reliability of scientific conclusions from cognitive modeling depends critically on the reliability of parameter estimation, yet estimating parameters remains a universal challenge particularly when data are too limited to constrain them. In such cases, multiple sets of parameters may explain the experimental data equally well within the same model, raising the question of which parameters are scientifically meaningful. We refer to this problem as *parameter ambiguity*. In this paper, we investigate parameter ambiguity in reinforcement learning under two optimization methods. We employ the de facto Nelder-Mead method (`fminsearch`) and a neural network trained to estimate parameters using a modern deep learning pipeline, which has seen limited application in cognitive modeling. Across ten decision-making datasets, we consistently find that the two methods produce substantially different parameter estimates despite achieving nearly identical fitting performance. To address this ambiguity, we introduce a systematic evaluation framework that goes beyond predictive accuracy to assess generalizability, robustness, identifiability, and test-retest reliability, thereby offering principled guidance on which parameter estimates should inform scientific inference. Applying this framework reveals that the neural network with a deep learning pipeline outperforms across these metrics. Our study establishes parameter ambiguity as an underappreciated challenge with significant implications for scientific replicability, highlighting that the choice of optimization method is a critical factor shaping scientific conclusions. We advocate for our multi-faceted evaluation approach to ensure reliable scientific inference and for broader integration of modern deep learning pipelines into cognitive modeling.

## 1 Introduction

Cognitive models are widely used in neuroscience and psychology to investigate the cognitive processes underlying behavior in humans and animals. These processes are described by model parameters that characterize how agents learn from experience, make decisions, and adapt to their environments. Reinforcement learning (RL) is one of the most successful frameworks for modeling decision-making as value learning from rewards and task-related feedback (Rescorla and Wagner, 1972; Sutton and Barto, 1998). Neuroscientific studies have identified neural substrates that implement RL computations (Lee et al., 2012; Niv, 2009; Schultz et al., 1997), supporting their biological plausibility.

RL models typically rely on a small number of free parameters to capture the cognitive processes driving behavior. These parameters are not merely tunable quantities used to fit models; they represent interpretable psychological constructs (Wilson and Collins, 2019), such as learning rates for value updating and inverse temperatures reflecting decision noise. Accurate and reliable parameter estimation is therefore essential for linking these quantities to cognitive processes and neural mechanisms, and for drawing meaningful scientific inferences (Browning et al., 2015; Huys et al., 2021; Suthaharan et al., 2021).

In practice, however, reliable estimation is often hindered by constraints common in human and animal experiments, such as limited data, behavioral variability, model complexity, and task designs with low diagnostic power for isolating specific cognitive mechanisms. These challenges can give rise to the problem of *parameter ambiguity*: different combinations of parameters explain the observed behavior equally well. In such cases, the estimated parameters may be influenced by the inductive biases of the optimization method — an important factor but often treated as a minor implementation detail in cognitive modeling.

In this study, we empirically demonstrate the prevalence of parameter ambiguity in RL models across *ten* diverse value-based decision-making datasets by comparing two optimization methods. The first is the Nelder-Mead simplex method (Nelder and Mead, 1965), which serves as the default in MATLAB’s `fminsearch` (The MathWorks, Inc., 2024) and SciPy’s `fmin` (Virtanen et al., 2020). Its off-the-shelf availability has contributed to its widespread adoption in cognitive modeling (for representative applications, see Akam et al., 2015; Busemeyer and Myung, 1987; Busemeyer and Diederich, 2010; Dowman et al., 2016; Forstmann et al., 2011; Gershman, 2018; Li et al., 2023; Pouget et al., 2011; Shinn et al., 2020; Wilson et al., 2019; Yechiam and Busemeyer, 2005). The second method involves training a neural network to estimate RL parameters within a modern deep learning pipeline. While the optimization pipeline used in deep learning have achieved considerable success in other domains, they are rarely used in cognitive modeling (but see Giallanza et al., 2024).

Surprisingly, both methods often yield markedly different parameter distributions across subjects in all datasets, despite achieving similar predictive performance on held-out data. This highlights the universality of parameter ambiguity and suggests that predictive performance alone is insufficient to justify the validity of estimated parameters. It raises a critical question: which parameters should be used to draw scientific inferences?

To address this parameter ambiguity, we propose a systematic evaluation framework that assesses parameter reliability along four key dimensions: generalizability, quantified by the gap between training and test loss; robustness, measured by sensitivity to parameter perturbations; identifiability, evaluated through parameter recovery from simulated data; and test-retest reliability in longitudinal datasets. We find that the neural network achieves superior performance across all four dimensions. This multi-faceted approach provides a principled basis for determining which parameter estimates are scientifically meaningful. Our analysis underscores both the necessity of comprehensive evaluation and the potential of modern deep learning pipelines as a robust alternative for parameter estimation in cognitive modeling.

## 2 Results

### 2.1 Datasets and models

To investigate the problem of parameter ambiguity in cognitive modeling, we selected ten diverse decision-making datasets involving humans and animals performing bandit tasks with varying action spaces, reward structures, and time horizons (for details see Sec. 4.1). We denote each dataset by the study name and year, such as `Wilson2014`). We fit all datasets using the same widely adopted formulation of the RL model, in which the action value *Q*_*t*_ for the chosen action *A*_*t*_ at trial *t* is updated as follows:

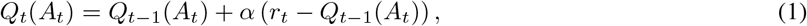

where *r*_*t*_ is the reward at trial *t*, and *α* denotes the learning rate specifying how quickly the values are updated. For action selection, we use a Softmax policy:

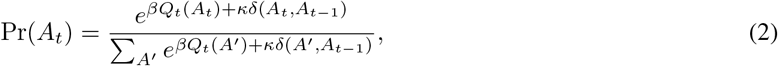

where *β* is the inverse temperature specifying the action determinism, *κ* is the choice perseverance parameter reflecting an inertia tendency, and *δ*(*A*_*t*_, *A*_*t−*1_) equals 1 if *A*_*t*_ = *A*_*t−*1_ and 0 otherwise.

### 2.2 Parameter ambiguity under different optimization choices

We now investigate parameter ambiguity under optimization choices in practice across ten datasets. We apply the Nelder-Mead method and a neural network with a modern deep learning pipeline (hereafter referred to as *neural network* for simplicity) to fit RL models to behavioral data by estimating the RL parameters (*α, β, κ*) for each subject and experimental condition (Fig. 1a). The Nelder-Mead method directly optimizes the RL parameters using 64 grid-based initializations (see Sec. 4.3 for details). In contrast, the deep learning pipeline (Fig. 1b) uses a single-layer neural network that takes subject- and condition-level embeddings as input and outputs the corresponding RL parameters (see Sec. 4.2 for details). The network parameters are optimized with the gradient-based optimizer Adam (Kingma and Ba, 2017), using weight decay (Krogh and Hertz, 1991; Loshchilov and Hutter, 2019) and early stopping (Amari et al., 1997; Finnoff et al., 1993). The performance of both optimization methods is assessed through cross-validation: parameters are optimized on training data, and performance is measured on held-out test data (see Sec. 4.3 for details).

**Figure 1:**
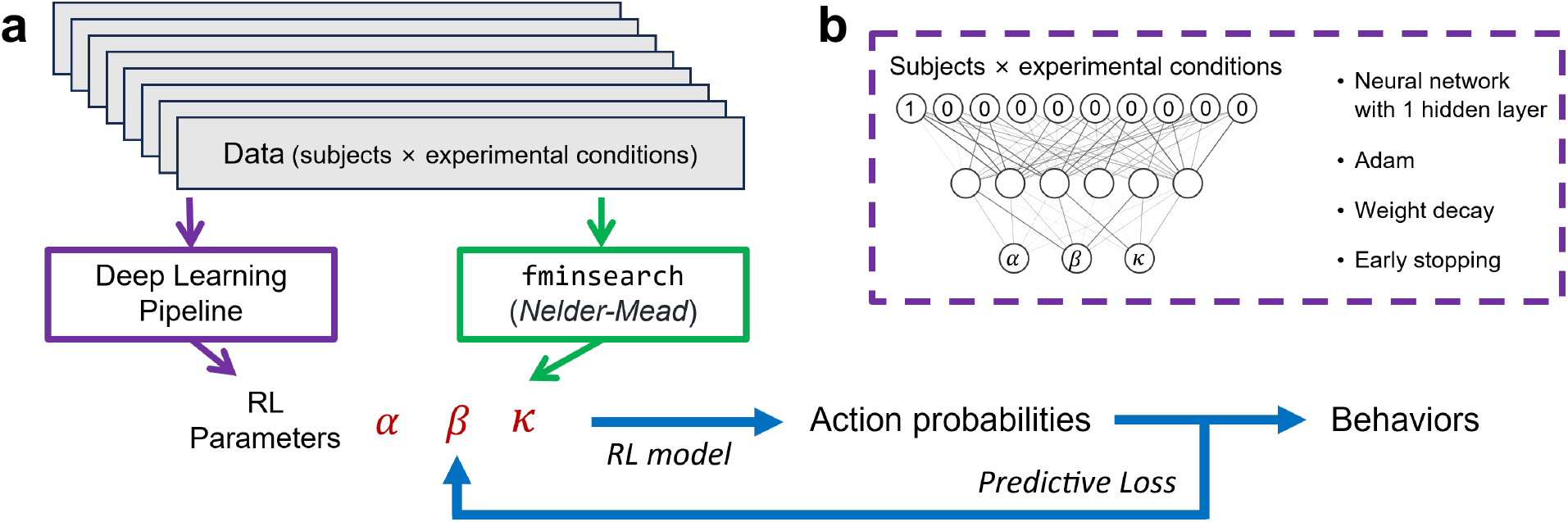
Illustration of the data fitting procedure. **a**, Both methods optimize parameters in RL models for each subject and experimental condition to predict choice behavior. **b**, A one-layer neural network combined with a deep learning pipeline that includes the AdamW optimizer, weight decay, and early stopping to estimate RL parameters.

Both methods perform similarly on held-out test data across ten datasets (Fig. 2a). No significant difference in average performance is observed (Fig. 2b, Wilcoxon signed-rank test: *W* = 12.0, *p* =.131, Cohen’s *d* = 0.031). To examine whether the two methods yield similar parameter estimates when predictive performance is similar, we show the joint distributions of parameters across subjects and experimental conditions in ten datasets. Distinct RL parameter distributions are consistently observed across ten datasets (Fig. 3; see Fig. A.1 for marginal distributions). While the distributions of *κ* overlap more across datasets, the joint distributions involving *α* and *β* differ markedly. And we find the neural network tends to generate fewer extreme estimates (e.g., *α* in `Steingroever2015` and *β* in `Dubois2022`). Moreover, the covariance structure of RL parameters diverges substantially between the two methods (see also parameter correlations in Fig. A.2). As a result, cognitive interpretations based on parameter relationships, such as the coupling between learning rate and exploration, may vary depending on the optimization method.

**Figure 2:**
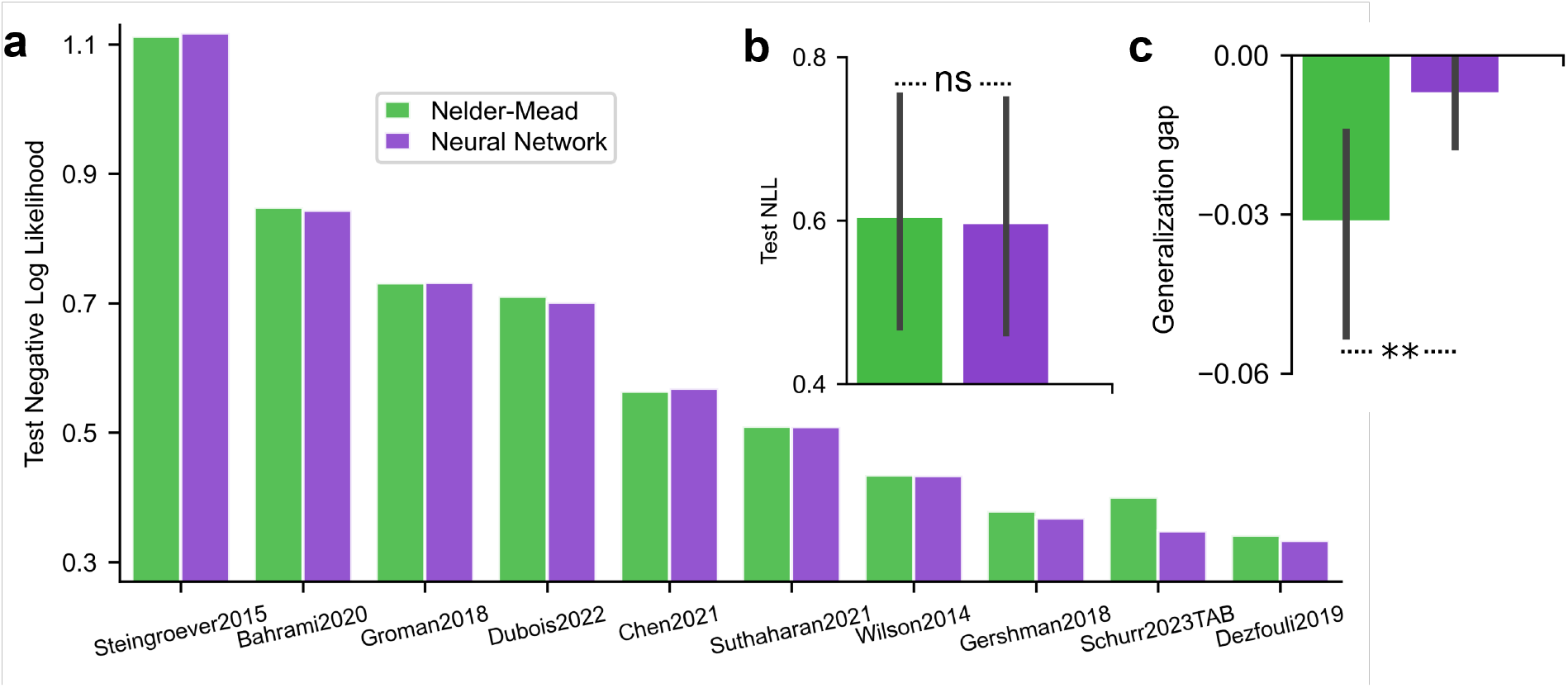
Comparison of Nelder-Mead and neural network methods across ten decision-making datasets shows equivalent fitting performance. **a**, Predictive performance of RL models optimized with each method. The neural network achieves a negative log-likelihood (lower is better) comparable to that of the Nelder-Mead method on held-out test data. **b**, Average predictive performance across all datasets. Both methods yield similar negative log-likelihoods on held-out test data. **c**, Generalization gap, defined as the difference in negative log-likelihood between training and test data, averaged across datasets. The Nelder-Mead method shows a larger generalization gap, suggesting potential overfitting to the training data. Error bars in **b**,**c** denote 95% confidence intervals across the ten datasets. ns: not significant. **: *p* < 0.01.

**Figure 3:**
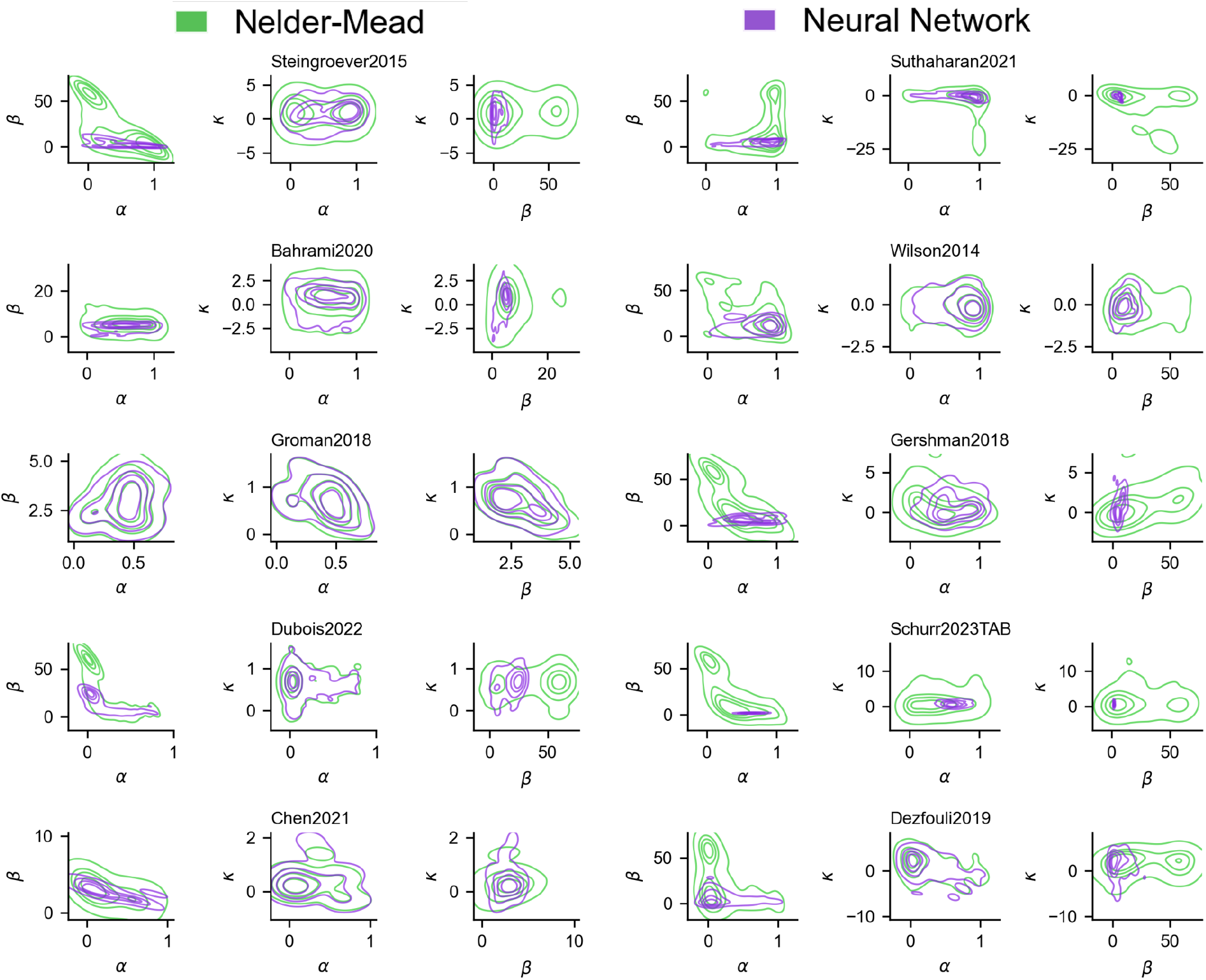
The Nelder-Mead method and neural network yield distinct two-dimensional joint distributions of RL parameters. Each subplot presents a kernel density estimate of the joint distribution between pairs of RL parameters (*α, β, κ*), aggregated across all subjects and experimental conditions within a dataset. Contour lines indicate density levels, with separate colors denoting each estimation method.

Beyond distributional differences, parameter ambiguity may distort relative comparisons across subject groups, such as whether one subject has a higher learning rate than another, which is central to scientific inference. Therefore, an important test of parameter ambiguity is whether the rank ordering of subjects’ parameters is preserved across optimization methods. For both methods, we ranked subjects by each RL parameter within each dataset and evaluated agreement using Kendall’s *τ*(Fig. 4; see also Pearson’s *r* in Fig. A.3). Rank agreement across methods was substantially lower for *α* and *β*, particularly in datasets with smaller sample sizes. As sample size increased, the inter-subject parameter rank structure became more consistent across methods. For *κ*, rank correlation remained high and stable across datasets, suggesting that its estimated parameter rank structure is robust to the choice of optimization method. Overall, the two methods produced different rank orderings of the estimated RL parameters across subjects. This result highlights that parameter ambiguity can distort conclusions about individual differences, a challenge that becomes especially pronounced as sample size decreases.

**Figure 4:**
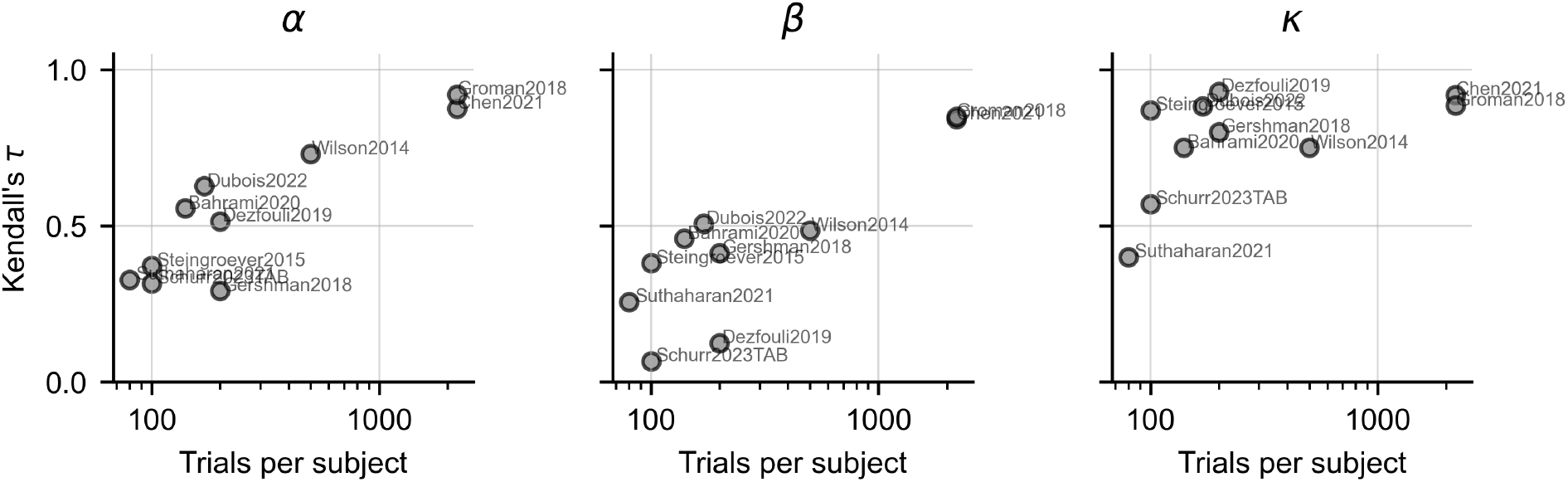
Agreement in subject-wise rank ordering of RL parameters across optimization methods. Each panel shows Kendall’s *τ* rank correlation between the Nelder-Mead method and the neural network for each RL parameter (*α, β, κ*) as a function of the average number of trials per subject.

The finding that two methods produce different RL parameter estimates despite equivalent predictive performance across ten datasets demonstrates the prevalence of parameter ambiguity and raises concerns about the reliability of scientific interpretations. Without additional metrics to distinguish among estimates, conclusions based on a single value may be ambiguous or misleading. In the remainder of our analysis, we introduce a systematic framework that goes beyond predictive performance to address this ambiguity.

### 2.3 Assessing generalization via train-test performance gap

The first dimension of our evaluation framework is generalizability. For estimated RL parameters to be interpretable as reflecting a subject’s underlying cognitive processes, they must generalize to new situations rather than apply only to the data on which they were fitted. Otherwise, the estimates merely capture noisy patterns specific to the training trials, a problem known as overfitting. We assess the risk of overfitting for both methods by comparing the generalization gap (Keskar et al., 2017), defined as the difference in negative log-likelihood between training and test sets. The neural network exhibits a smaller generalization gap than the Nelder-Mead method (Fig. 2c, Wilcoxon signed-rank test: *W* = 0.0, *p* =.002, Cohen’s *d* = *−*0.956), indicating reduced risk of overfitting and greater generalizability.

### 2.4 Parameter sensitivity and robustness analysis

Robust interpretation of estimated cognitive parameters requires that small perturbations do not substantially alter model performance. This robustness reflects the “flatness” of the identified minima in which the parameters are estimated. Flat minima are associated with improved generalization (Hochreiter and Schmidhuber, 1997; Neyshabur et al., 2017; Xie et al., 2021). To quantify this, we perturb the RL parameters estimated by both methods on each trial by adding independently and identically distributed (i.i.d.) zero-mean Gaussian noise with varying variance. We then compute the loss (negative log-likelihood) between subjects’ behavior and the predictions of RL models using the perturbed parameters. The resulting loss increases are averaged across subjects and experimental conditions within each dataset (Fig. 5a), and then across datasets (Fig. 5b). Across all datasets, loss increases with greater noise magnitude, indicating reduced performance. Notably, the loss increases more slowly for the neural network than for the Nelder-Mead method (Wilcoxon signed-rank test: *W* = 49.0, *p* <.001), suggesting reduced sensitivity to parameter noise and thus greater robustness, potentially reflecting flatter parameter minima.

**Figure 5:**
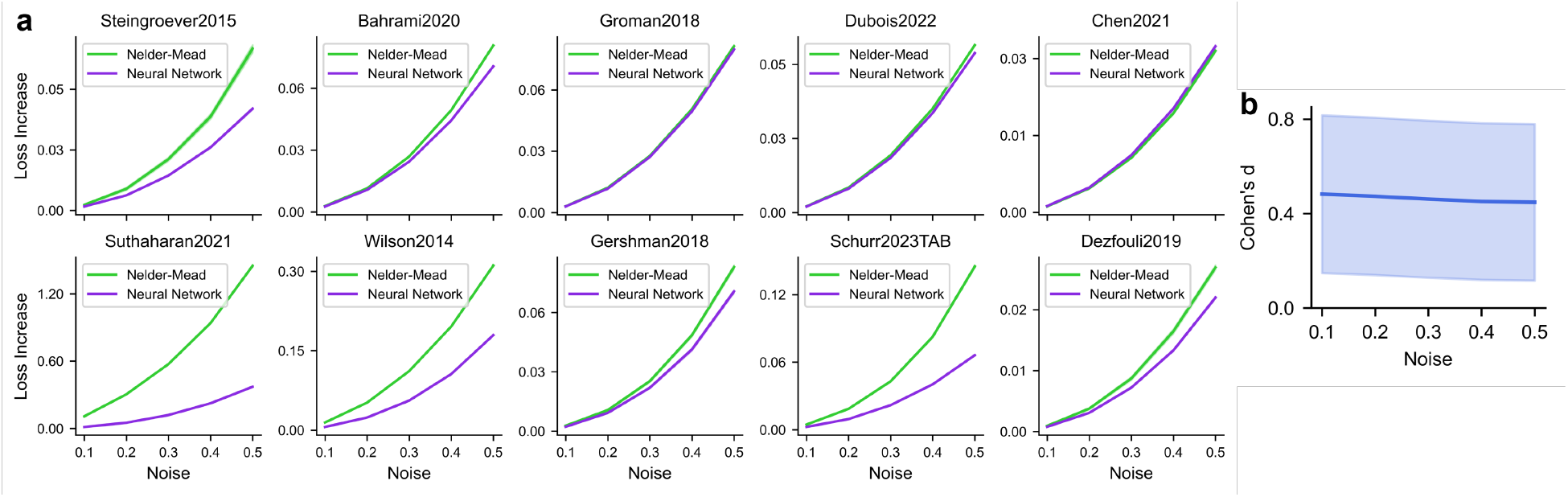
Model robustness under parameter perturbations at varying noise levels. **a**, Parameters estimated by the neural network exhibit smaller increases in loss in response to injected white Gaussian noise, compared to the Nelder-Mead method, meaning lower expected sharpness and greater robustness to perturbation. The shaded area (mostly invisible) represents the 95% confidence interval across 10,000 simulations. **b**, Effect size of the difference in loss increase between the Nelder-Mead method and the neural network, in which the Nelder-Mead method shows a greater loss increase across ten datasets. The shaded area denotes the 95% confidence interval across ten datasets.

### 2.5 Evaluating parameter identifiability using parameter recovery

An important and commonly used criterion for meaningful parameter interpretation is identifiability, often measured by the ability to reliably recover ground-truth parameters from simulated data (Wilson and Collins, 2019). If ground-truth parameters cannot be recovered, estimates derived from empirical data may lack interpretability with respect to underlying cognitive processes. Identifiability evaluates whether a task and model provide sufficient information to determine the parameters of interest. Here, we extend this framework to address parameter ambiguity arising from optimization choices. When different optimization methods yield distinct parameter estimates but achieve equivalent fits for the same task and model, we consider the estimates from the method with superior recovery performance to be more scientifically meaningful.

We compare parameter recovery across four representative tasks that span diverse task structures (for the rationale behind task selection and implementation details, see Sec. 4.5). We generate simulated data using 125 parameter combinations, with five values spanning a plausible range for each parameter, and then re-estimate those parameters by fitting the same model structure to the data while treating the RL parameters as free. We assess recovery performance using the correlation and mean absolute error between estimated and ground-truth parameters. For both methods, recovery improves as the number of trials per subject increases, converging to similar levels with sufficient data (3200 trials). However, residual errors between estimated and ground-truth parameters remain in some tasks even with sufficient trials (e.g., Fig. 6b; 2-armed instrumental learning, third row), suggesting that identifiability may be fundamentally constrained by the task and model structure.

**Figure 6:**
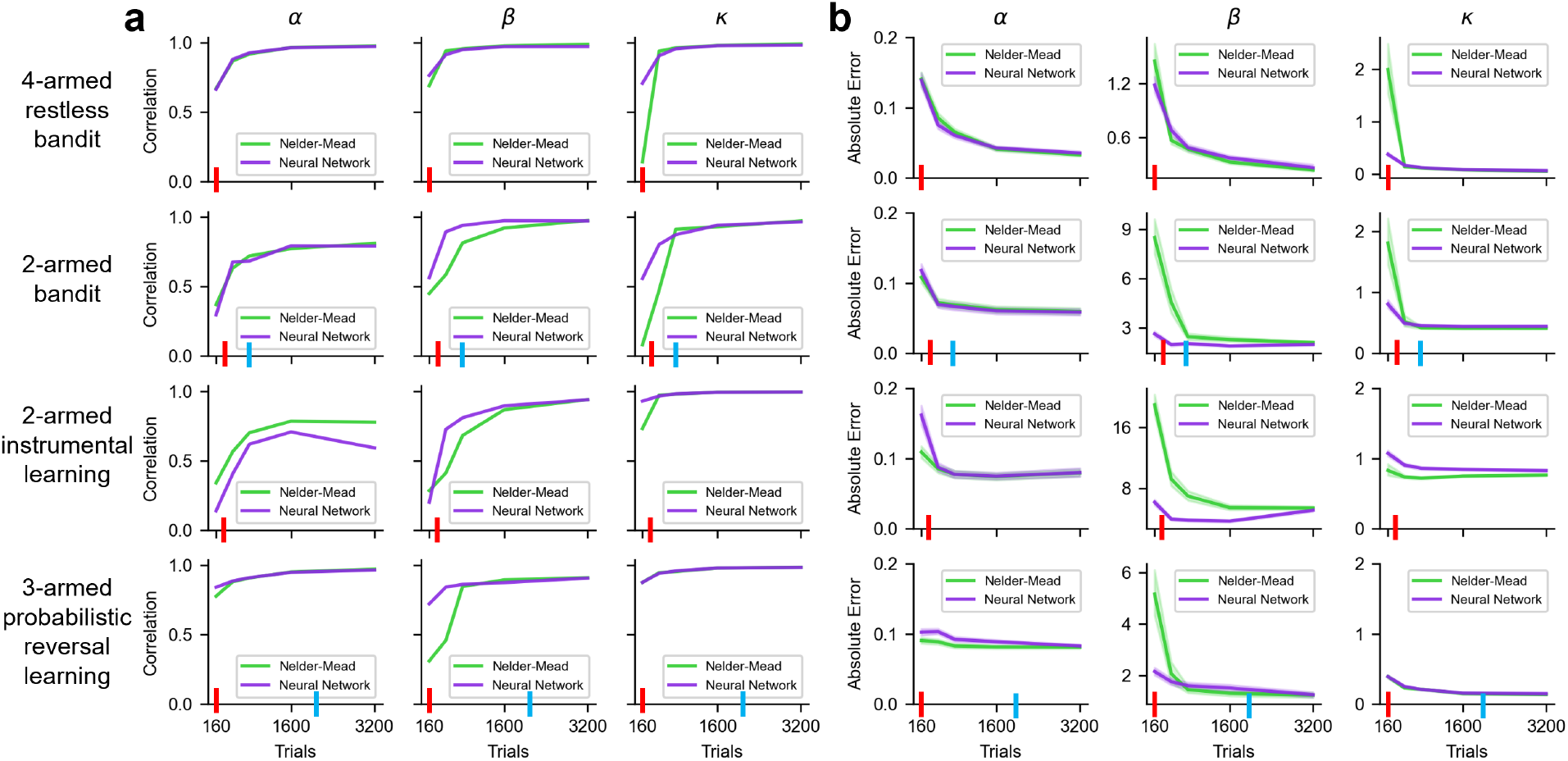
Parameter recovery using the Nelder-Mead method and the neural network across varying amounts of simulated data from an RL model. The neural network achieves higher or comparable **(a)** correlations with ground-truth parameters and **(b)** lower or comparable absolute errors across most tasks. The shaded region (barely visible) denotes the 95% confidence interval across all parameter combinations. Red and blue ticks on the x-axis indicate the average number of trials per subject in our datasets: 4-armed restless bandit (red, `Bahrami2020`); 2-armed bandit (red, `Gershman2018, Schurr2023TAB`; blue, `Wilson2014`); 2-armed instrumental learning (red, `Dezfouli2019`); and 3-armed probabilistic reversal learning (red, `Suthaharan2021`; blue, `Groman2018`).

The neural network outperforms the Nelder-Mead method in most tasks when the number of trials per subject is limited, a range comparable to typical sample sizes in psychological experiments (red and blue bars in Fig. 6). This result is notable given the common assumption that deep learning methods require large datasets to be effective. Our findings indicate that the neural network is particularly well-suited for typical dataset sizes in computational modeling, as it more accurately recovers ground-truth parameters in this regime.

### 2.6 Test-retest reliability of estimated parameters

Finally, we evaluate test-retest reliability, which provides a direct and intuitive link to scientific inference. High reliability of parameter estimates is especially valuable for computational phenotyping (Patzelt et al., 2018), as it supports the prediction of clinical outcomes and the development of treatments. Test-retest reliability assesses the consistency of scores for the same subjects tested under similar conditions at different time points, making it laborious to obtain. We analyze a unique dataset, `Schurr2023TAB` (Schurr et al., 2024), in which subjects performed a two-armed bandit task over 12 consecutive weeks, enabling us to measure correlations across repeated assessments for the same individuals. Following the exclusion criteria and methodology of the original study (Schurr et al., 2024), we compute the intra-class correlation (ICC) of the RL model parameters, a widely used measure of test-retest reliability.

We also control for differences in fitting performance between the two methods by selecting a subset of subjects (41 out of 90) for whom the predictive loss, measured as negative log-likelihood (NLL), differs by less than 0.02 (Fig. 7a; NLL quantiles: Q1 = -0.003, Q3 = 0.012. Our results remain consistent when applying different threshold values). After controlling for fitting performance, the two methods produce distinct parameter distributions (Fig. 7b). The neural network also demonstrates greater robustness to parameter perturbations (Fig. 7c). Crucially, the neural network yields significantly higher ICC values for the estimated RL parameters (Fig. 7d): *β*: *t*(40) = 3.30, *p* =.002, Cohen’s *d* = 0.52; *κ*: *t*(40) = 4.88, *p* <.001, Cohen’s *d* = 0.95. The only exception is *α*, which shows no significant difference (*t*(40) = 0.71, *p* =.482, Cohen’s *d* = 0.06), indicating stronger test-retest reliability overall. These findings suggest that the neural network produces more consistent parameter estimates across repeated measurements from the same individuals.

**Figure 7:**
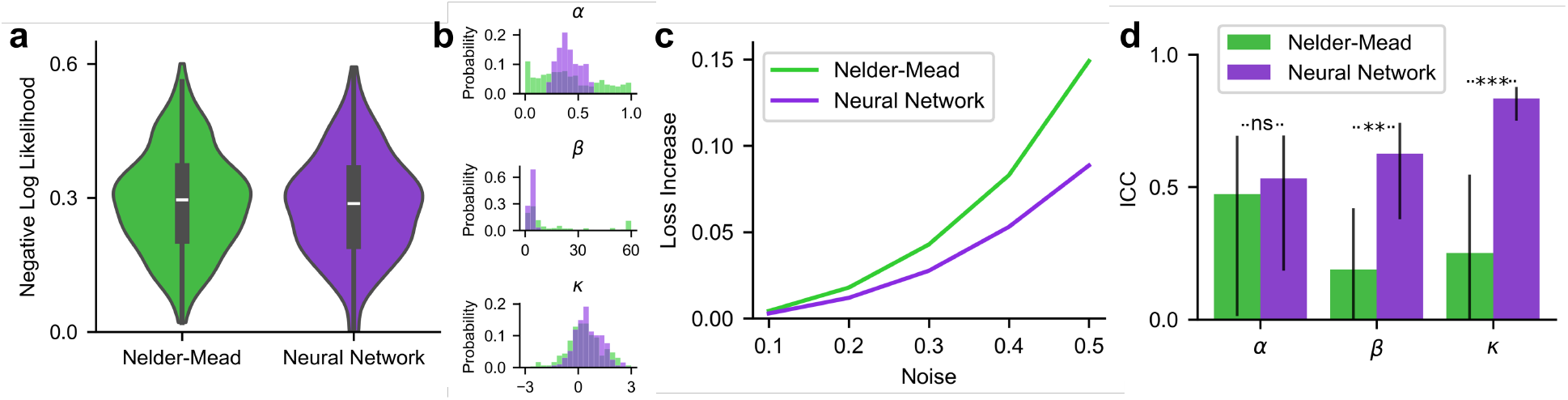
Test-retest reliability of the Nelder-Mead method and neural network on the Schurr2023TAB dataset with multiple measurements from the same subjects. A subset of subjects was selected to control for fitting performance (see main text). **a**, Both methods show similar distributions of fitting performance. **b**, The two methods yield distinct parameter distributions. **c**, The neural network exhibits greater robustness to parameter perturbations. The shaded area (mostly not visible) represents the 95% confidence interval across 10,000 simulations. **d**, The neural network achieves higher ICC values for the estimated parameters across eight weekly measurements from the same subjects, compared to the Nelder-Mead method. Error bars represent 95% confidence intervals across subjects. Significance levels are denoted as follows: ns = not significant, * *p* <.05, ** *p* <.01, *** *p* <.001.

## 3 Discussion

Parameters in cognitive models serve as interpretable constructs for describing cognitive processes and drawing scientific inferences. Replicable inference depends on accurate and reliable parameter estimation. In this paper, we demonstrate the prevalence of *parameter ambiguity*, a critical yet underappreciated vulnerability in cognitive modeling, where different parameter estimates explain the data equally well within the same model. We reveal this problem by comparing optimization choices across ten datasets, showing different joint parameter distribution and inter-subject ranking order, presenting a serious challenge to the interpretability and replicability of scientific findings based on cognitive modeling. If two labs use the same model on similar data but different optimizers, they could arrive at different scientific conclusions without realizing the source of the discrepancy.

To address this ambiguity, we propose a systematic evaluation framework that assesses estimated parameters in terms of generalizability, robustness, identifiability, and test-retest reliability. Our framework provides principled guidance on which RL parameter estimates should be used for inference. To our knowledge, this is the first systematic evaluation of how optimization methods influence parameter estimation across multiple metrics and datasets in cognitive modeling.

We also show that a neural network with modern optimization techniques performs better on our evaluation framework than the commonly used Nelder-Mead method, the default optimizer in MATLAB’s `fminsearch` and SciPy’s `fmin`.

In cognitive modeling, constraints on dataset size particularly with human participants, the type of task can be employed, the intrinsic variability of behavior, and other factors can all contribute to a non-convex fitting loss landscape. Such a landscape allows multiple parameter sets to explain the data equally well. Consequently, the inductive bias of the chosen optimization method determines the estimated parameters. To address this issue, we propose four evaluation metrics to identify which parameter sets are most suitable for scientific inference.

We first evaluate generalizability. Better parameter estimates should capture core behavioral patterns that extend to unseen data rather than overfitting noise in the training data. We quantify this using the generalization gap, defined as the difference in fitting loss between training and test sets, which measures the risk of overfitting.

We also evaluate robustness. Reliable parameter estimates should be insensitive to small perturbations, meaning slight changes do not substantially alter the model’s behavior. We formalize it as the sensitivity of the loss to parameter perturbations, which is rarely applied in cognitive modeling (but see Frank, 2014). Lower sensitivity indicates greater flatness, which is linked to improved generalization through “wide flat minima” (Fig. 8), a well-established concept in the machine learning literature (Hochreiter and Schmidhuber, 1997; Dinh et al., 2017; Hoffer et al., 2017; Neyshabur, 2017; Xie et al., 2021).

**Figure 8:**
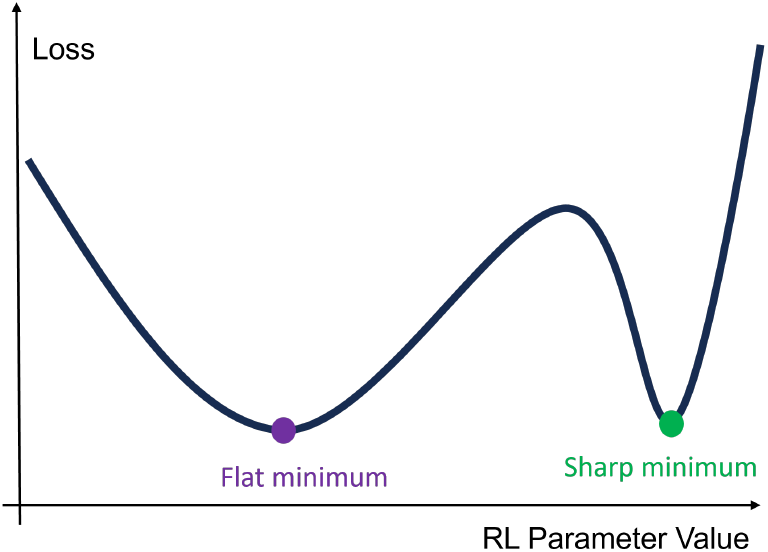
Illustration of parameter flatness and sharpness. In a non-convex loss landscape, different parameter estimates may produce similar losses. flatter minima, where loss remains stabler under small parameter perturbations, are generally considered better solutions and are often associated with stronger generalizability.

The statistical power of inference also depends on the identifiability of cognitive parameters. We use parameter recovery to assess identifiability by testing whether ground-truth parameters can be accurately recovered from simulated data generated with them. High identifiability indicates that parameter estimates have greater power and are more likely to reflect the underlying generative processes. Identifiability is closely related to the Fisher information, which underlies the theoretical consistency of maximum likelihood estimation. We found that the neural network aligned more closely with ground-truth parameters than the Nelder-Mead method across four widely used decision-making tasks, especially in the low-data regime typical of human behavioral experiments.

Identifiability and robustness may seem contradictory at first glance. Identifiability requires high sensitivity of the loss function to parameter changes: once parameters deviate from their true values, the loss increases sharply. In contrast, robustness requires low sensitivity to small perturbations in parameter values. However, the two properties are complementary. Identifiability concerns recovery of the ground truth, whereas robustness ensures that the loss landscape remains relatively flat around the solution, providing stability against perturbations (see Appendix A.2 for details). An effective cognitive model and corresponding experiment balance both properties, such that higher Fisher information ensures identifiability, while a bounded Hessian ensures robustness.

Our final metric, test-retest reliability, is the most intuitive and a key factor for scientific inference, as it reflects the consistency of parameter estimates across repeated measurements. Previous research shows that the test-retest reliability of RL parameters is generally low to moderate (Schaaf et al., 2024; Schurr et al., 2024). We find that the neural network achieves significantly higher test-retest reliability, enabling more accurate and robust characterization of cognitive processes. This improvement further supports the use of cognitive parameters as stable biomarkers, with applications in computational psychiatry and cognitive phenotyping, including longitudinal studies of individual differences in personality and development (Huys et al., 2021; Montague et al., 2012; Patzelt et al., 2018).

To demonstrate the prevalence of parameter ambiguity under optimization choices, we use a single formulation of the RL model, one 10-armed bandit task, and two optimizers. Future studies could extend this approach to different RL model formulations, a variety of tasks, other cognitive models, and additional optimizers.

Our study focused on point-estimation methods, which are commonly used in practice. Our framework can be equally applied to Bayesian point estimates (e.g., MAP) or posterior summaries. A systematic comparison with full Bayesian inference represents an important direction for future research, as these methods provide posterior uncertainty but also introduce challenges such as prior specification and convergence diagnostics, which lie beyond the present scope.

While our primary goal is to demonstrate parameter ambiguity and provide a framework for addressing it, the finding that the neural network within a deep learning pipeline performs better across four metrics is also noteworthy. We conduct ablation analyses (Appendix A.3) to investigate the sources of this advantage. The results indicate that shared representations modestly enhance identifiability, particularly in low-data regimes, while other components, including the overparameterized neural network, the Adam optimizer, weight decay, and early stopping, are also critical for accurate model fitting. For example, stochastic gradient-based optimizers such as Adam tend to bias solutions toward flatter minima that is more generalizable (Chaudhari et al., 2019; Xie et al., 2021). Nevertheless, no single factor fully accounts for the observed advantages, and a complete mechanistic explanation remains beyond the scope of this work.

In conclusion, our study highlights the prevalence of parameter ambiguity in cognitive modeling and identifies optimization choice as a critical yet often overlooked factor influencing cognitive parameters. We advocate for systematic evaluation of cognitive parameters to ensure reliable scientific inference and to strengthen their links to psychological constructs and neural mechanisms. We also show that the neural network with a modern deep learning pipeline produces more reliable parameter estimates, making it a promising and scalable tool for cognitive modeling.

## 4 Method

### 4.1 Datasets

The ten datasets include several types of decision-making tasks performed by both humans and animals, featuring different levels of uncertainty, various reward structures, and horizon lengths.

`Wilson2014` is a two-armed horizon task (Wilson et al., 2014) that involves two options (arms). The value difference between the two arms is sampled from 4, 8, 12, 20, and 30, and rewards are drawn from a normal distribution. The task includes a forced-choice phase in the first four trials, which controls the uncertainty level regarding the value of each option. In the free-choice phase, the horizon is set to either 1 or 6, allowing researchers to study how decision-making changes with the amount of available information in the future. This dataset includes data from multiple labs and experiments (Wilson et al., 2014; Feng et al., 2021; Somerville et al., 2017; Waltz et al., 2020; Sadeghiyeh et al., 2020; Zajkowski et al., 2017), with *n*_subjects_ = 609 and *n*_trials_ = 754,890.

`Dubois2022` is a three-armed version of this horizon task (Dubois and Hauser, 2022). This version increases task complexity by introducing a third arm, requiring more strategic considerations when choosing the best option. This dataset includes *n*_subjects_ = 1,012 and *n*_trials_ = 173,120.

`Chen2021` is a two-armed restless bandit task in mice (Chen et al., 2021) that uses binary rewards. In this task, the reward probability for each arm changes gradually over time, requiring the mice to continuously adapt their choices based on these shifting probabilities. This dataset includes *n*_subjects_ = 32 and *n*_trials_ = 70,778.

`Bahrami2020` is a four-armed restless bandit task in humans (Bahrami and Navajas, 2020). Arm values range from 0 to 100 and follow an Ornstein-Uhlenbeck process toward 50. Initial values are sampled from a normal distribution, and rewards are drawn based on the arm’s current value. The task therefore incorporates both reward stochasticity and environmental volatility due to the fluctuating arm values. This dataset includes *n*_subjects_ = 965 and *n*_trials_ = 139,816.

`Suthaharan2021` is a three-armed probabilistic reversal learning task in humans (Suthaharan et al., 2021). Subjects choose between three arms, each associated with different reward probabilities. In the first two blocks, probabilities are set to [0.9, 0.5, 0.1], and in later blocks, they shift to [0.8, 0.5, 0.2]. After ten consecutive selections of the best option, the reward probabilities are reversed, allowing researchers to examine how individuals respond to changing reward contingencies. This dataset includes *n*_subjects_ = 1,012 and *n*_trials_ = 173,120.

`Groman2018` is a three-armed probabilistic reversal learning task conducted in mice (Groman et al., 2018). Reward probabilities are [0.8, 0.5, 0.2], enabling cross-species comparisons of decision-making strategies. This dataset includes *n*_subjects_ = 17 and *n*_trials_ = 38,475.

`Gershman2018` is a two-armed bandit task in humans (Gershman, 2018) with a horizon length of 10 and mean reward values sampled from a normal distribution. In experimental condition 1, the mean reward for one arm is fixed at 0, while the other arm’s mean reward varies. In experimental condition 2, both arms have varying mean reward values, allowing researchers to examine how subjects balance exploration and exploitation. This dataset includes *n*_subjects_ = 89 and *n*_trials_ = 17,800.

`Schurr2024TAB` replicates experimental condition 2 of the two-armed bandit task to assess test-retest reliability (Schurr et al., 2024). Subjects engage in the two-armed bandit task over 12 consecutive weeks. The dataset includes *n*_subjects_ = 90 and *n*_trials_ = 78,820. Participant exclusion criteria follow those reported in the original paper.

`Dezfouli2019` is a two-armed instrumental learning task in humans (Dezfouli et al., 2019). Rewards are binary, with one arm offering higher reward probabilities [0.25, 0.125, 0.08] compared to the other arm with a reward probability of 0.05. The duration of each experimental condition is fixed, enabling analysis of learning behavior over a defined period, particularly how subjects adapt to consistent reward probabilities. This dataset includes *n*_subjects_ = 101 and *n*_trials_ = 132,251.

`Steingroever2015` is the Iowa Gambling Task (Steingroever et al., 2015), in which subjects choose from four decks of cards, each associated with different reward and punishment structures. This task assesses decision-making under risk, as subjects must learn which decks are more beneficial in the long term based on experienced rewards and penalties. This dataset includes data from multiple labs and experiments, with *n*_subjects_ = 577 and *n*_trials_ = 62,525.

### 4.2 Neural network model

The neural network we used in our deep learning pipeline consists of a single hidden layer with no activation functions and is defined as follows:

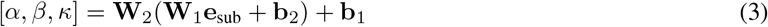

where **e**_sub_ ∈ ℝ^*n×n*^ is a one-hot embedding for each subject per experimental condition, with *n* representing the total number of subjects multiplied by the number of experimental conditions in a given task. The weight matrix **W**_1_ ∈ ℝ^*h×n*^ maps the one-hot embedding to a hidden representation, while **b**_1_ ∈ ℝ^*h*^ is the corresponding bias term. The second weight matrix **W**_2_ ∈ ℝ^3*×h*^ maps the hidden representation to the RL parameters, with bias term **b**_2_ ∈ ℝ^3^. While this two-layer linear network is equivalent to a single-layer network in its representational power, they have substantially different training dynamics (Saxe et al., 2013). The output RL parameters *α, β, κ* are then fed in the RL models.

In PyTorch implementation, the entire RL recurrent computation and neural network feedforward computation are both embedded within the computational graph, enabling gradients to be computed via backpropagation through time.

### 4.3 Model fitting

The optimization objective is the negative log-likelihood of the observed choice data:

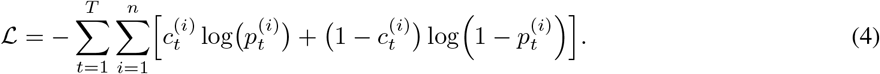

Here, 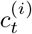 denotes the actual choice of subjects *i* on trial *t* and 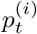 denotes the model’s prediction probability for the subjects’ choices (Eq. 2).

#### 4.3.1 Cross validation

To mitigate overfitting and ensure a fair comparison between both optimization methods, we implemented a rigorous cross-validation protocol. For the Nelder-Mead method, we employed five-fold cross-validation, allocating 80% of the data for training and 20% for testing. For the neural network, we adopted a nested cross-validation approach (Ji-An et al., 2023). Specifically, 70% of the data was designated for training, 12.5% for validation (used for early stopping and hyperparameter tuning), and 17.5% for testing. The outer loop comprised eight folds, while the inner loop included five folds.

Given the variability in datasets, experimental designs, and trial counts, we tailored the cross-validation data splits accordingly. For datasets comprising multiple short blocks with identical trial counts (Gershman, 2018; Schurr et al., 2024), we partition data into different blocks. For datasets with multiple short blocks of varying trial counts (each block containing fewer than 40 trials) (Wilson et al., 2014; Dubois and Hauser, 2022), we divided the data into non-overlapping 40-trial segments. When necessary, we added padding to maintain structural uniformity; padded trials were excluded from loss computation. For datasets with relatively few blocks and fewer than 200 trials per block (Bahrami and Navajas, 2020; Suthaharan et al., 2021; Reed et al., 2020; Steingroever et al., 2015), we split the data into non-overlapping trial sets by applying different masks for training, validation, and testing (Hans et al., 2024). For datasets with few blocks but more than 200 trials (Chen et al., 2021; Groman et al., 2018), we divided each session into non-overlapping blocks of 100 trials and add padding when necessary.

#### 4.3.2 Hyperparameter

For the Nelder-Mead optimization method, parameter fitting for each subject within a given experimental condition underwent 64 independent initializations. For RL parameter estimation, we constructed a grid of initial points by evenly sampling four values for each parameter: *α ∈* [0, 1], *β ∈* [0, 60], and *κ ∈* [*−* 5, 5]. Our choice of 64 grid-based initializations represents a rigorous and computationally intensive procedure, designed to provide a strong baseline for the Nelder-Mead method and exceeding the common practice in many studies.

To enhance the robustness of the deep learning pipeline, we augmented the training data by adding Gaussian noise *N* (0, 0.01) to continuous rewards. Early stopping was applied when validation performance did not improve after 50 training epochs, ensuring retention of models with the lowest validation loss. This strategy effectively prevented overfitting. Hyperparameters were optimized over the following search space: learning rate selected from {0.01, 0.02, 0.03}, and hidden layer size *h* from {30, 50, 100}, weight decay from {0.01, 0.1}. Other optimizer hyperparameterswere set to the default valu s in Pytorch for AdamW: betas=(0.9, 0.999), eps=1e-08. Constraints were enforced on RL parameter estimation, ensuring *α ∈* [0, 1] and *β ≥* 0.

The best model was selected based on the lowest negative log-likelihood on the test data, averaged across all folds. For the RL parameters, we averaged them over cross-validation folds. In the Nelder-Mead method, we used all five folds. In the deep learning pipeline, we use five selected folds (outer *j* and inner fold 0, where *j* = 0, 1, 2, 3, 4). Our results remained consistent for other fold selection protocols.

### 4.4 Estimating parameter robustness

We assessed parameter robustness by perturbing model parameters and measuring the resulting change in the loss function. Specifically, we added white Gaussian noise at each trial to the fitted parameters 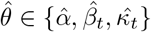 estimated by each optimization method. The noise was independently sampled from a zero-mean Gaussian distribution, with variance scaled by the magnitude of the subject-specific parameter:

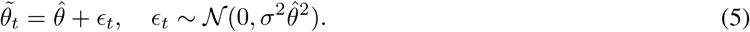

This formulation ensures that the perturbation scale is proportional to the fitted parameter for each subject, allowing robustness to be assessed relative to each parameter’s magnitude. The noise variance coefficient *σ*^2^ was selected from *{*0.1, 0.2, 0.3, 0.4, 0.5 *}*. For each predefined *σ* and optimization method, we generated 10,000 perturbed parameter sets. Each set was used to simulate model predictions, and the corresponding loss was computed as the negative log-likelihood between perturbed model output probabilities and observed behavioral data. We defined robustness (expected sharpness) as the average increase in loss, quantified by the difference between the perturbed and original losses.

### 4.5 Parameter recovery

To capture the range of behavioral patterns observed across the ten datasets, we selected four representative tasks for parameter recovery. Specifically, the 4-armed restless bandit task is based on `Bahrami2020`, using the same generative process but without controlling for identical reward schedules; the original task included only three fixed schedules. The 2-armed bandit task corresponds to those used in `Gershman2018` and `Schurr2024TAB`, and is structurally similar to `Wilson2014`. However, uncertainty manipulations and forced-choice trials in the Horizon task are not directly relevant to the model-free RL framework used in our analysis. The 2-armed instrumental learning task is adapted from `Dezfouli2019`, restricted to the [0.25, 0.08] reward condition. Finally, the 3-armed probabilistic reversal learning task is based on `Suthaharan2021` and `Groman2018`, using a reward structure of [0.9, 0.5, 0.1]. We excluded `Steingroever2015` because the Iowa Gambling Task variants use arbitrarily defined reward contingencies that lack a clear generative structure. We did not include `Chen2021`, another restless bandit task, as its design closely resembles other restless bandit tasks. Lastly, we omitted `Dubois2022` because of ambiguity in the task structure described in the original paper.

For each task, we selected simulated parameters within the most plausible range to perform the parameter recovery test (Wilson and Collins, 2019). This range was approximately defined using the interquartile range (second-to-third quartile) of parameter distributions obtained from our two methods. For tasks where subjects exhibited highly concentrated parameter distributions, we slightly expanded the range. This resulted in the following parameter ranges for each task: *α ∈* [0.3, 0.9], *β ∈* [4, 10], and *κ ∈* [0, 1.5] for the four-armed restless bandit task (Bahrami and Navajas, 2020); *α ∈* [0.3, 0.8], *β ∈* [4, 15], and *κ ∈* [*−*0.5, 1.5] for the two-armed bandit task (Gershman, 2018; Schurr et al., 2024); *α ∈* [0.05, 0.35], *β ∈* [1, 20], and *κ ∈* [0, 3] for the two-armed instrumental learning task (Dezfouli et al., 2019); and *α ∈* [0.4, 0.9], *β ∈* [4, 12], and *κ* [*∈ −*1.5, 0.5] for the three-armed probabilistic reversal learning task (Suthaharan et al., 2021; Reed et al., 2020; Groman et al., 2018).

Within these ranges, we uniformly sampled five values for each parameter. This sampling effectively decouples correlations between the ground-truth parameters. Using these 125 parameter combinations, we simulated data for 1, 3, 5, 10, and 20 sequence, with each sequence consisting of 160 trials. This resulted in datasets containing 160, 480, 800, 1,600, and 3,200 trials per parameter set.

For the four-armed restless bandit task and the three-armed probabilistic reversal learning task, each sequence contained a single block of 160 trials. In the three-armed probabilistic reversal learning task, we used reward probabilities of [0.8, 0.5, 0.2]. In the two-armed bandit task, each sequence consisted of eight blocks of 20 trials, and we used experimental condition 2 from (Gershman, 2018). In the two-armed instrumental learning task, each sequence consisted of two blocks of 80 trials, and we used reward probabilities of [0.125, 0.05], corresponding to the condition in which one arm had a higher reward probability. These choices were made to align with the original task designs.

We then estimated the parameters from the simulated data using both optimization methods across different data sizes, following the same model fitting and selection procedure described in Methods Model fitting. Due to the computational cost of model recovery and the fact that we had 125 distinct parameter combinations, we performed the recovery procedure once for each combination. While this introduces some stochasticity at the level of individual simulations, the large number of combinations provides a robust basis for assessing overall trends in recovery performance.

### 4.6 Parameter test-retest reliability

To assess the test-retest reliability of RL parameters over multiple measurements, we computed non-parametric bootstrapped intraclass correlations, ICC(2,1), using 1,000 samples, following methods from (Enkavi et al., 2019; Vallat, 2018). For each RL parameter, we used the mean of the posterior distribution as a point estimate and calculated the ICC based on the data of 90 participants with 12 sessions each. However, ICC cannot be computed when data is missing. To address this, we included all participants but restricted the analysis to the maximal number of sessions with complete data, using eight weekly measurements from the same subjects.

## Acknowledgments

We thank all the researchers who generously shared their datasets, making the comparative analysis in this study possible. This work was primarily conducted while HDX and RCW were at the University of Arizona and was supported by National Institutes of Health grant R01AG061888 awarded to RCW. We also acknowledge the use of the High Performance Computing (HPC) facilities at the University of Arizona, which provided essential computational resources for this research.

## Author contributions

Hua-Dong Xiong, Conceptualization, Formal analysis, Data curation, Investigation, Methodology, Project administration, Software, Visualization, Writing - original draft, Writing - review and editing, Conceived the work; Li Ji-An, Conceptualization, Methodology, Project administration, Writing - original draft, Writing - review and editing, Conceived the work; Marcelo G. Mattar, Supervision, Writing - review and editing; Robert C. Wilson, Resources, Supervision, Funding acquisition, Writing - review and editing.

## A Appendix

### A.1 Supplementary figures

**Figure A.1:**
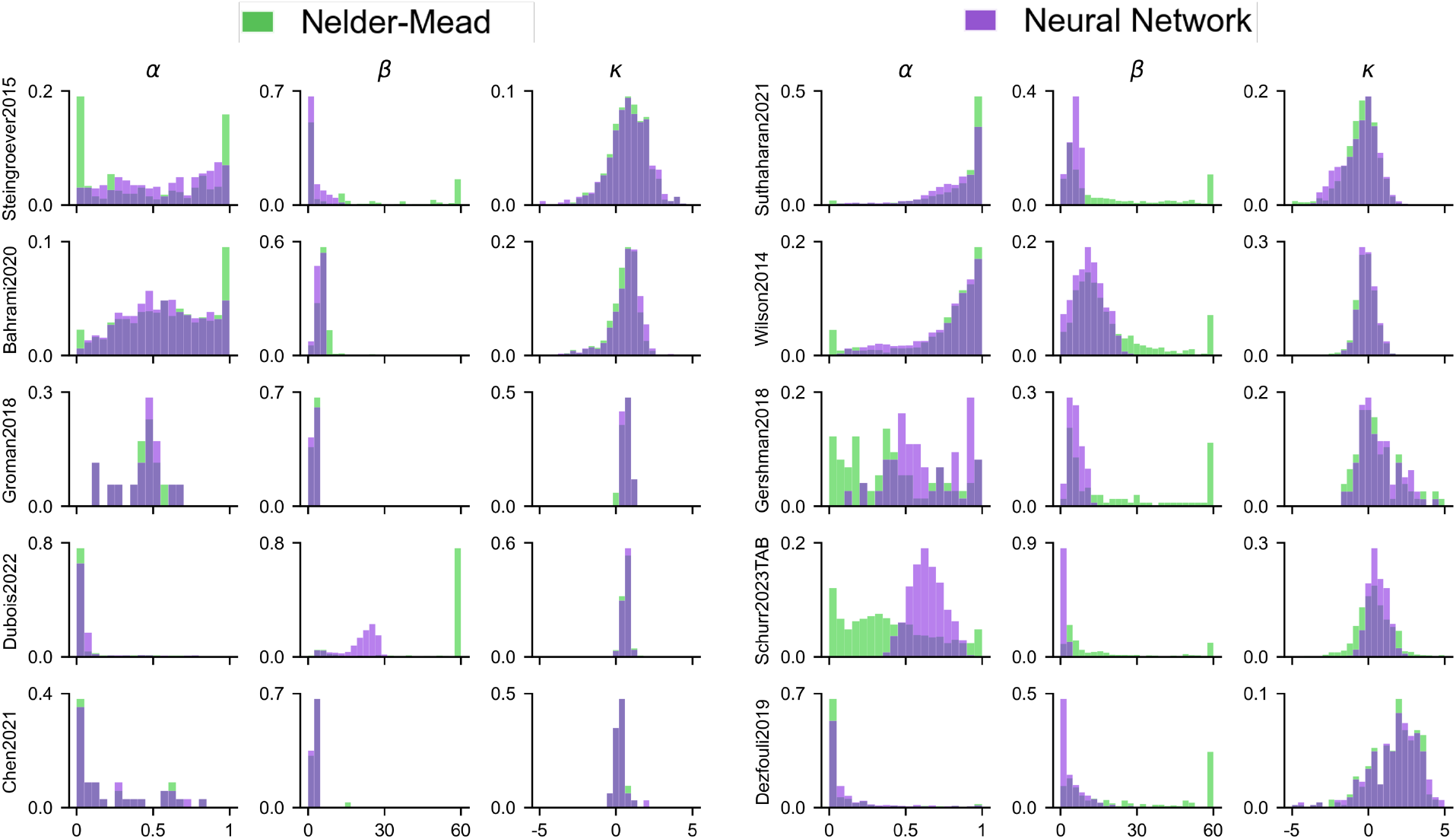
Marginal distributions of RL parameters estimated by the Nelder-Mead and deep learning pipeline. Each panel depicts the distribution of a single parameter (*α, β, κ*) across all subjects and experimental conditions within a dataset. The two methods produce distinct parameter distributions across ten datasets.

**Figure A.2:**
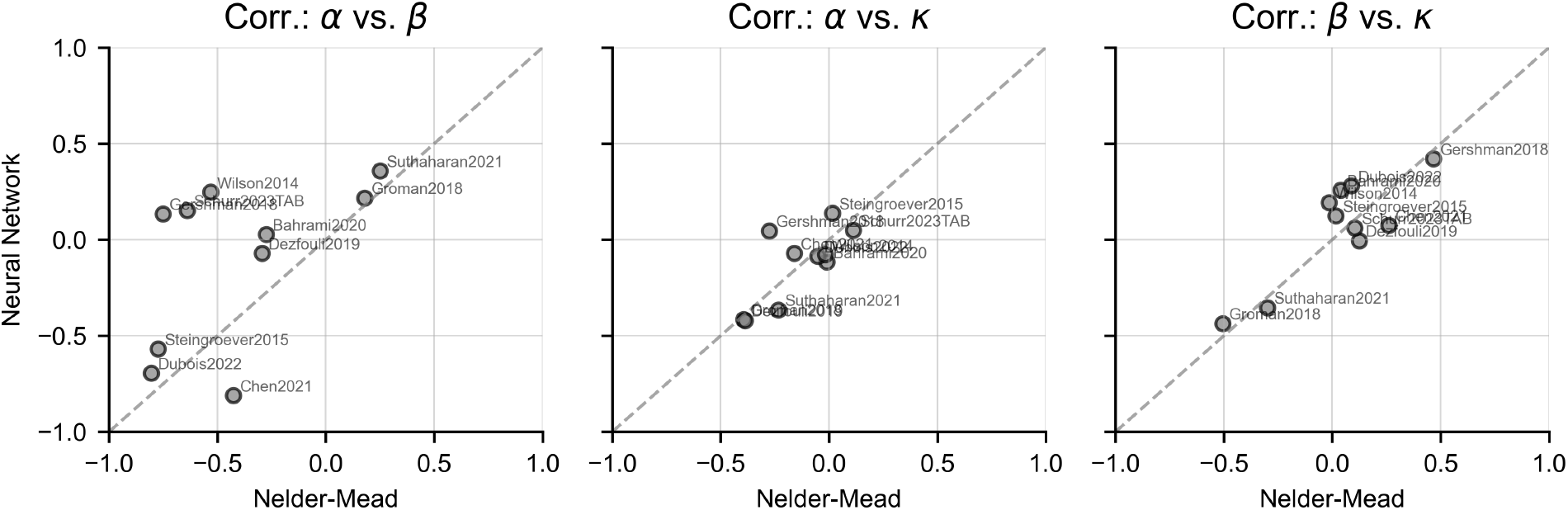
Parameter correlations estimated by the Nelder-Mead and deep learning pipeline. Each panel shows pairwise correlations among RL parameters (*α, β*, and *κ*), comparing estimates from the two optimization methods across datasets. Deviations from the identity line (i.e., *x* = *y*) indicate systematic shifts in the way parameters covary across individuals.

**Figure A.3:**
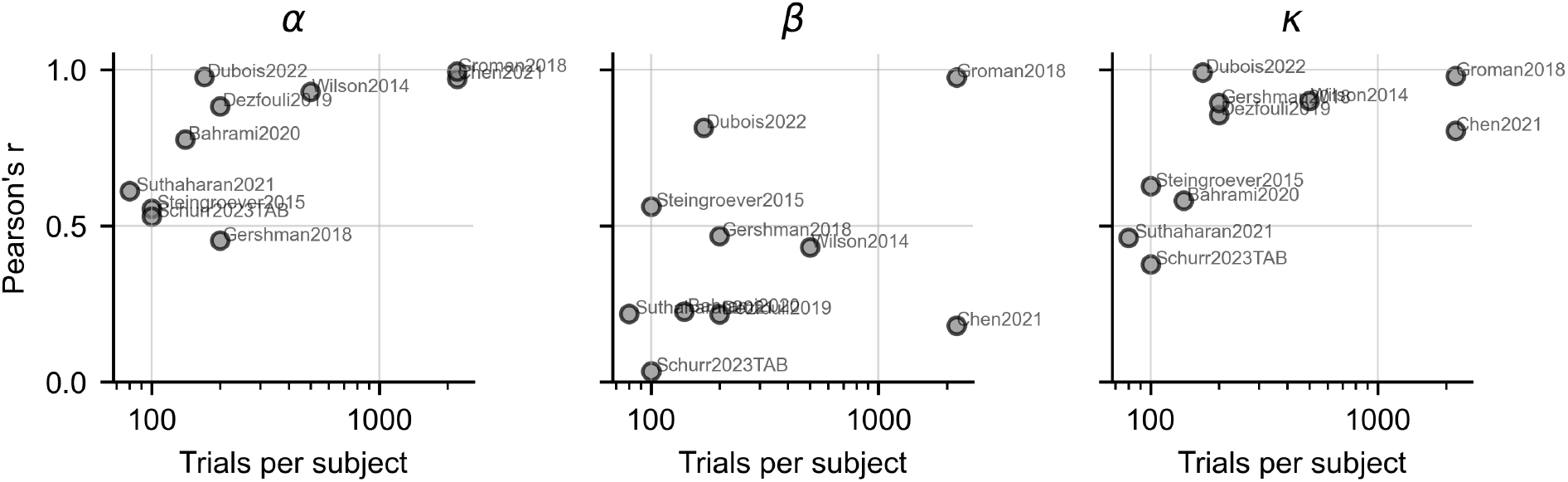
Agreement in subject-wise RL parameters across estimation methods. Each panel shows Pearson’s r correlation between the two estimation methods for a given RL parameter (*α, β*, or *κ*) as a function of sample size.

### A.2 Relations between identifiability, robustness, Hessian, and Fisher information

In this section, we examine the mathematical relationship between robustness, theoretically referred to as expected sharpness, and identifiability measured in this paper and well-known properties like Fisher information and the Hessian.

Below we consider a statistical model with one parameter *θ* (the generalization to more parameters is direct). The data *X* is generated by the model where the parameter takes *θ* = *θ*^*\**^ (*X ∼θ*^*\**^). The log-likelihood function (equivalent to the negative loss) is defined as

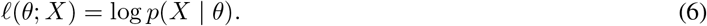

When *θ* is a local minimum, *l* can be locally expanded to the second-order approximation:

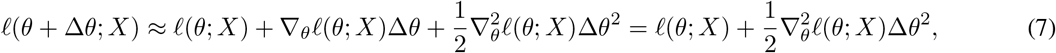

where the term *∇*_*θ*_*l*(*θ*; *X*) vanishes. The expected sharpness is then given by:

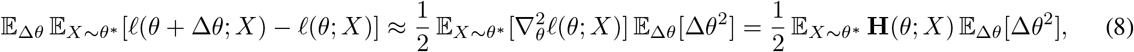

where **H** is the Hessian matrix which captures the local curvature. When Δ*θ* → 0, the equation between expected sharpness and expected Hessian becomes exact.

When *θ* = *θ*^*\**^, the Fisher information matrix, defined as

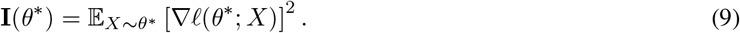

is equal to the negative expected Hessian (under standard regularity conditions):

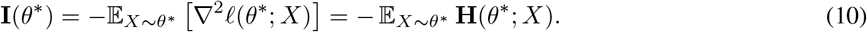

The Fisher information matrix measures the amount of information that the data *X* carries about the unknown ground-truth parameter *θ*^*\**^. Thus, it directly reflects the identifiability of the ground-truth parameters, as shown in the fact that the maximum likelihood estimator converges in distribution to a normal distribution:

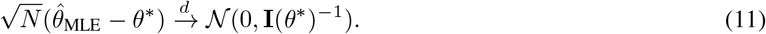

In practice, both the Fisher information matrix and the Hessian are estimated empirically:

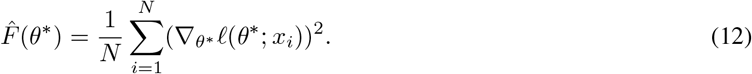

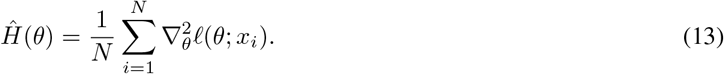

Specifically, these concepts measure distinct aspects of model parameters. The Fisher information for identifiability is always calculated at the ground truth parameters: if 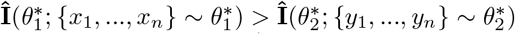, then 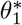 has higher identifiability. On the other hand, if 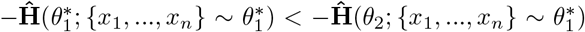, then then 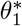 has lower expected sharpness (higher robustness) than *θ*_2_. Notice that *θ*_2_ is referring to a local minimum of the data generated by the ground-truth parameter 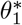. Therefore, 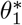 can have both high identifiability and high robustness.

### A.3 Shared representation learning helps better identifiability

Although we believe that all components of our deep learning pipeline—including the Adam optimizer, neural network architecture, early stopping, and weight decay—jointly contribute to its improved performance, we sought to isolate the specific contribution of shared representation. Our goal was to examine whether removing this component would yield similar predictive performance on unseen test data, but potentially reduce robustness, identifiability, or test-retest reliability.

We first confirmed that the use of Adam, early stopping, and weight decay are essential for achieving good fitting performance. Removing any of these components resulted in degraded performance on held-out data (data not shown), indicating their importance in generalization.

We then focused on the hypothesis about structure of neural network, that shared representation contributes to improved parameter estimation. The deep learning pipeline leverages shared structure across subjects and experimental conditions via shared representation, allowing the model to exploit similarities across individuals. In contrast, the Nelder-Mead method fits each subject and condition independently and cannot benefit from such shared information.

To isolate the role of shared representation, we ablated the neural network architecture by varying the number of hidden layers. When the network has zero hidden layers, it effectively implements a direct mapping from one-hot subject embeddings to RL parameters, preventing the model from learning shared representations. This ablation allows us to test whether performance benefits arise from shared representation or simply from the use of a neural network.

We found that the deep learning pipeline achieves similar test-set fitting performance regardless of the number of hidden layers (0, 1, or 2), as shown in Fig. A.4. This suggests that shared representation does not substantially affect predictive accuracy on held-out data.

**Figure A.4:**
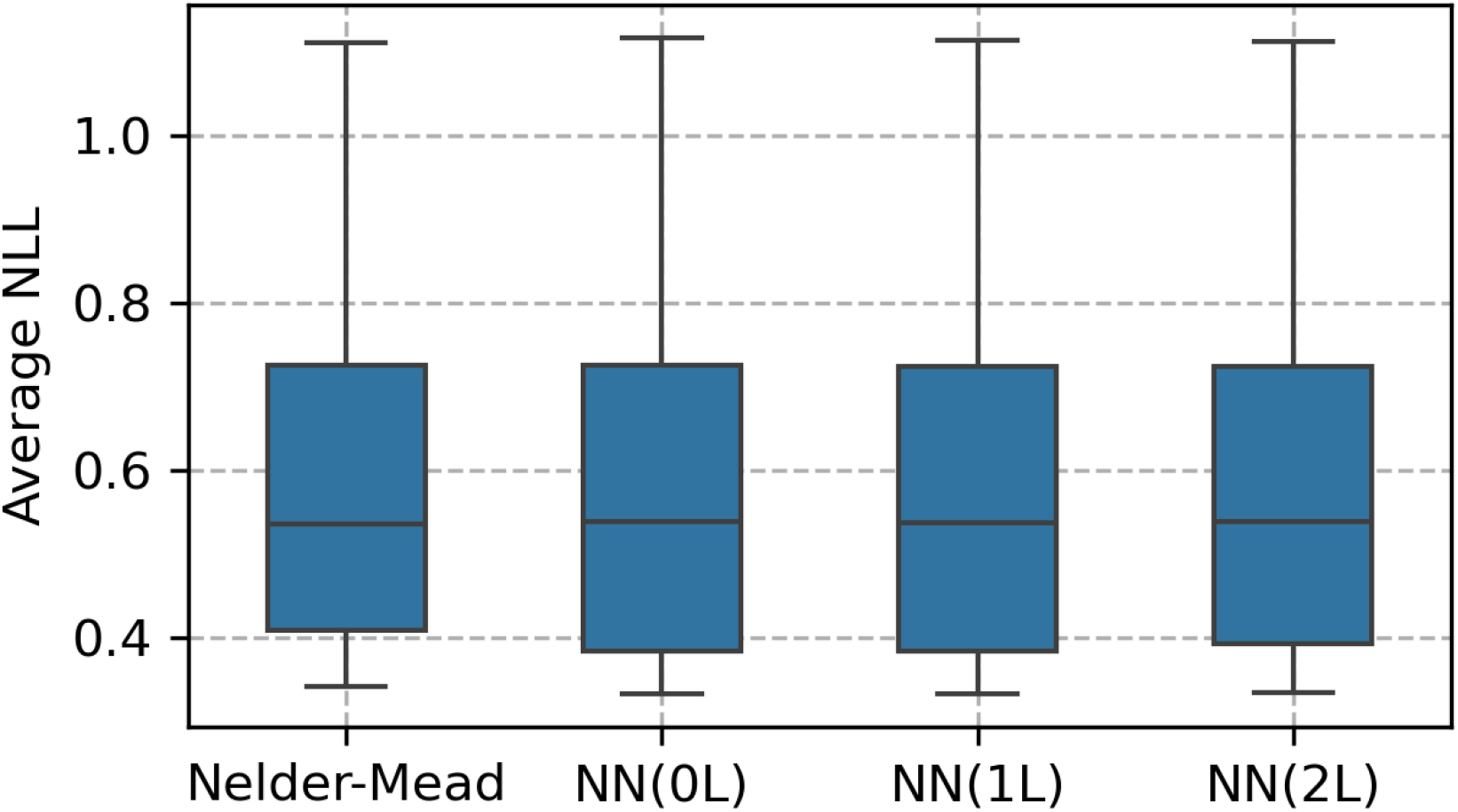
Average test-set fitting performance of the deep learning pipeline with 0 (0L), 1 (1L), or 2 (2L) hidden layers, compared to the Nelder-Mead method, across ten decision-making datasets.

We next examined robustness, measured as sensitivity to parameter perturbations. As shown in Fig. A.5, robustness was largely similar across architectures with 0, 1, and 2 hidden layers, suggesting that shared representation does not meaningfully improve robustness.

**Figure A.5:**
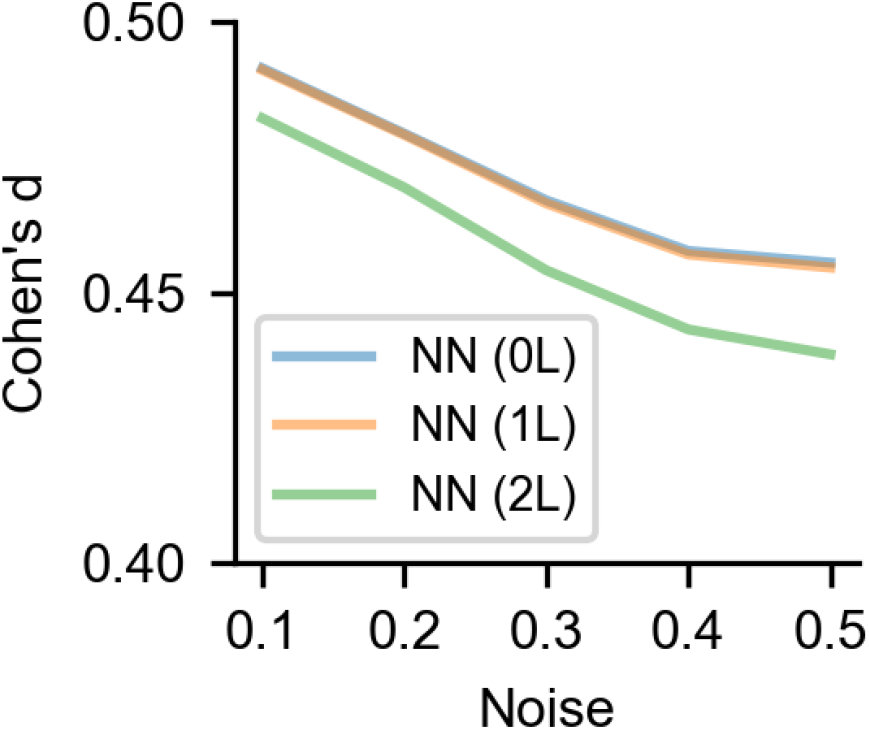
Effect size of the difference in loss increase under parameter perturbations for networks with 0 (0L), 1 (1L), or 2 (2L) hidden layers.

We also evaluated test-retest reliability on the `Schurr2023TAB` dataset. ICC values were largely consistent across network depths, though minor differences were observed in the *α* parameter. Overall, however, the pattern of results remained stable (Fig. A.6).

**Figure A.6:**
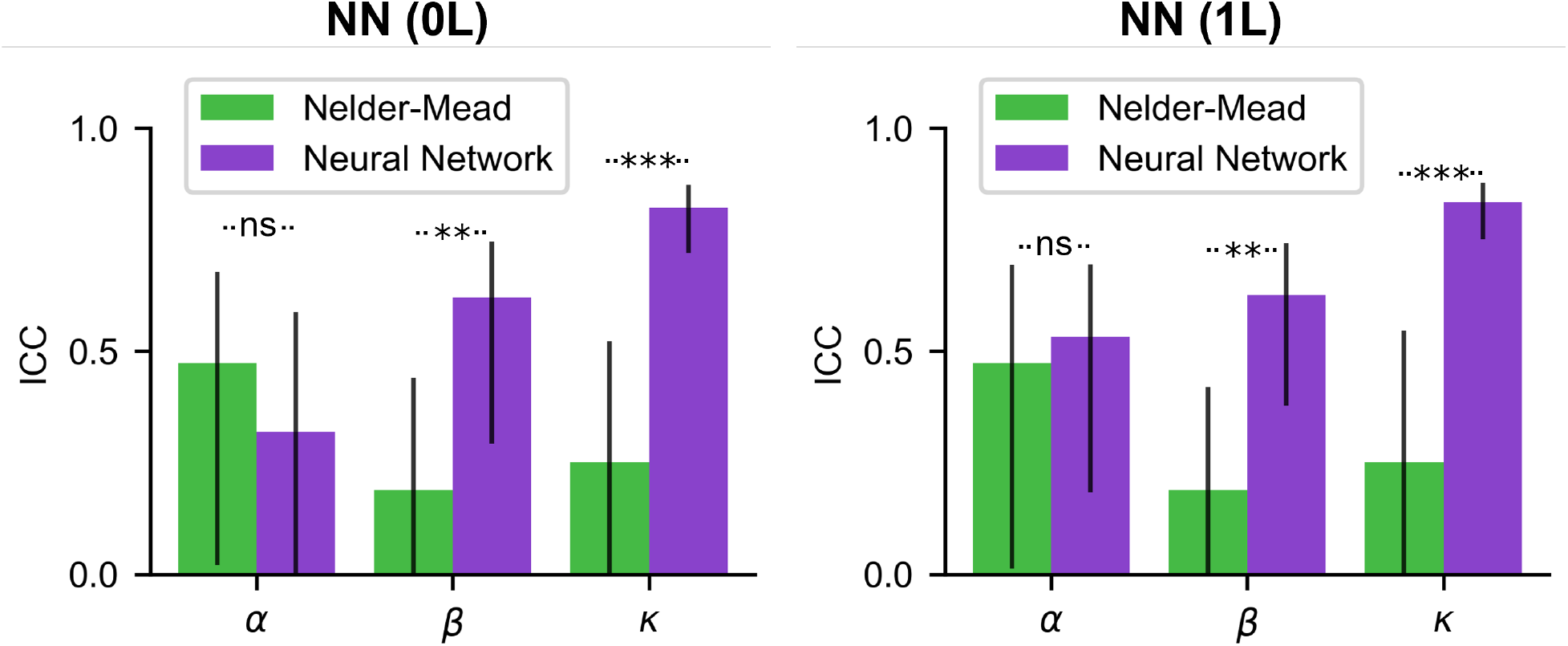
Test-retest reliability of the deep learning pipeline with 0 and 1 hidden layers show similar ICC values on the Schurr2023TAB dataset. Error bars represent 95% confidence intervals across subjects. Significance levels are denoted as follows: ns = not significant, * *p* <.05, ** *p* <.01, *** *p* <.001.

Importantly, we found that identifiability—measured by parameter recovery in simulated data—was reduced for the network with zero hidden layers, particularly for *β* and *κ* (Fig. A.7). This effect was most pronounced when trial counts per subject were limited. These results suggest that shared representation contributes to improved identifiability by enabling the model to leverage structural similarities across subjects, especially in data-limited regimes.

**Figure A.7:**
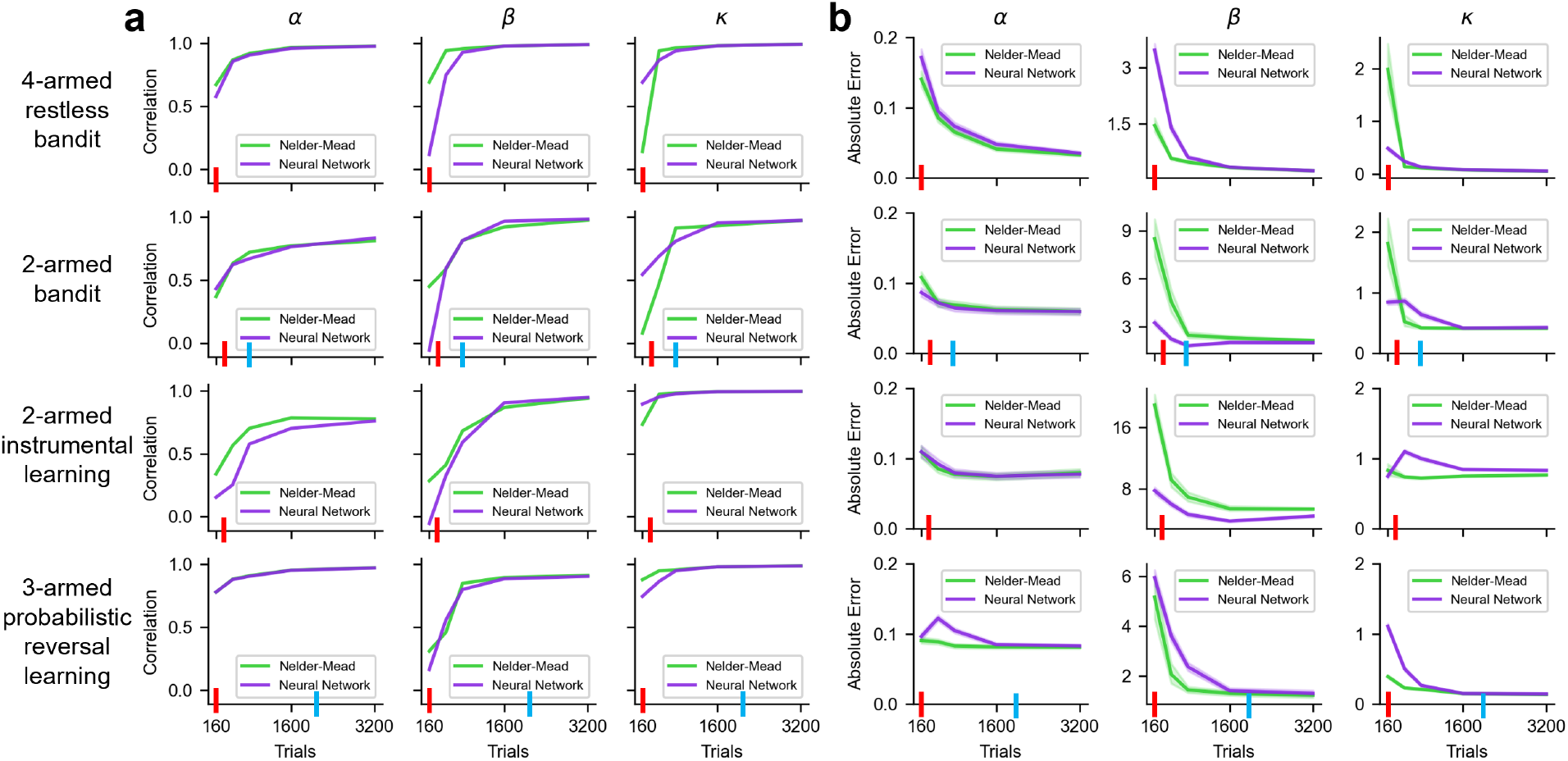
Parameter recovery with a neural network architecture using 0 hidden layers. Same analysis as Fig. 6, highlighting reduced identifiability without shared representation.

In summary, our ablation study indicates that while shared representation does not substantially affect predictive performance or robustness, it does improve identifiability, particularly in low-data settings. These findings underscore that the benefits of the deep learning pipeline emerge from the combination of its components, including modern optimizers, regularization techniques, and joint representations—none of which alone fully accounts for the observed improvements in parameter estimation.

